# Crystal structure and nanobodies against domain 3 of the malaria parasite fusogen *Plasmodium falciparum* HAP2

**DOI:** 10.64898/2025.12.01.691709

**Authors:** Frankie M. T. Lyons, Jill Chmielewski, Li-Jin Chan, Mikha Gabriela, Rainbow W. B. Chan, Amy Adair, Joshua Tong, Phillip Pymm, Kathleen Zeglinski, Quentin Gouil, Wai-Hong Tham, Melanie H. Dietrich

**Affiliations:** The Walter and Eliza Hall Institute of Medical Research, 1G Royal Parade, Parkville, Victoria 3052, Australia; Department of Medical Biology, The University of Melbourne, Parkville, Victoria 3010, Australia; Olivia Newton-John Cancer Research Institute; Heidelberg, Australia; School of Cancer Medicine, La Trobe University; Bundoora, Australia; Research School of Biology, The Australian National University, Canberra, ACT 2601, Australia

**Author notes:** Correspondence to: Melanie Dietrich and Wai-Hong Tham, The Walter and Eliza Hall Institute of Medical Research, 1G Royal Parade, Parkville, Victoria 3052, Australia, and.

**Keywords:** malaria, fusogen, transmission, crystallography, nanobody

## Abstract

Malaria parasites are transmitted to humans through a bite from an infected female *Anopheles* mosquito. Within the mosquito midgut, malaria parasite gametes are activated and undergo fertilisation. If parasite fertilisation is perturbed, this stops the transmission of malaria parasites from mosquito to human. One proposed target of transmission-blocking interventions is *Plasmodium falciparum* fusogen PfHAP2, which is essential for gamete fusion during parasite fertilisation. However, to date, no monoclonal antibodies or structures of PfHAP2 have been generated. We have identified nanobodies that bind specifically to domain 3 of PfHAP2 with nanomolar affinities, two of which show some cross-species reactivity with HAP2 of other *Plasmodium* species. The crystal structure of one nanobody in complex with domain 3 of PfHAP2 provides the first structural insights into this transmission-blocking target in *P. falciparum*.

## Introduction

Malaria is an ancient parasitic disease with global relevance today. *Plasmodium falciparum* is the deadliest species of malaria parasite, responsible for over 95% of malaria cases and approximately 600,000 deaths annually [1]. Transmission-blocking interventions aim to prevent the spread of disease by blocking parasite development inside the mosquito host. *P. falciparum* transmission begins when a female *Anopheles* mosquito bites an infected person and takes up blood containing the sexual stages of the malaria parasite, called gametocytes. Inside the mosquito midgut, male and female gametocytes emerge from red blood cells as gametes. Parasites undergo fertilisation whereby male microgametes fertilise female macrogametes. Fertilised parasites undergo several developmental stages in the mosquito, eventually generating thousands of infectious sporozoites that travel to the salivary glands, where they are poised to infect another person. Transmission from mosquito to humans can be stopped by blocking parasite fertilisation.

Parasite fertilisation requires attachment of male and female gametes followed by fusion of their membranes to form a zygote. Fusogens PfHAP2 and PfHAP2p are involved in membrane fusion [2]. HAP2 proteins belong to the fusexin superfamily, which includes fusogens responsible for critical biological process such as sexual reproduction, viral infection and syncytial development [3,4]. Other members of the superfamily include class II viral fusogens, such as protein E of tick-borne encephalitis virus, and the *Caenorhabditis elegans* somatic fusogen EFF-1, with which HAP2 shares structural and functional homology [5–8]. Fusexin1 is a structural homolog of HAP2 in archaea, the first fusexin to be described in prokaryotes [3].

HAP2 proteins are comprised of three domains (D1-D3), a stem region and a C-terminal transmembrane region. HAP2 is thought to exist as a monomer on the gamete surface, which trimerises during membrane fusion [5,9–11]. The prefusion structure of the algal species *Cyanidioschyzon merolae* reveals an extended, linear structure, with the three domains arranged as D3 – D1 – D2 in order of proximity to the transmembrane domain [10]. HAP2 interacts with the membrane of the opposite sex gamete via fusion loops in D2 and undergoes structural rearrangement during membrane fusion to pull the membranes together [5–7,9–12]. In the post-fusion crystal structure of *Chlamydomonas reinhardtii*, HAP2 shows a more compact structure in which D3 is folded back over D1 and D2 [5,11,12].

Work in the mouse malaria species *Plasmodium berghei* identified *P. berghei* HAP2 (PbHAP2) as having a critical role in membrane fusion. PbHAP2 is expressed on the surface of male gametocytes and gametes, and male knockout lines attach to female gametes but cannot progress to membrane fusion [13,14]. The structure of PbHAP2 D3 compared to algal and plant HAP2 structures shows structural conservation across species [8]. In *P. falciparum*, two HAP2 proteins have been characterised: PfHAP2 and PfHAP2p [2]. Both proteins are expressed in male gametocytes and microgametes, with PfHAP2p expression also detected in female stage V gametocytes [2]. Male knockout lines for both proteins are unable to form zygotes or infect mosquitoes, demonstrating non-redundant critical roles in male fertility for PfHAP2 and PfHAP2p [2]. Interestingly, a mammalian cell fusion assay indicated successful fusion required PfHAP2 and PfHAP2p to be expressed on both fusing cells, resembling the mechanism of action of *C. elegans* somatic fusogens [2]. The exact mechanism of action of gamete fusion in *P. falciparum* and the relationship between PfHAP2 and PfHAP2p is yet to be fully understood.

Polyclonal antibodies targeting HAP2 from *P. falciparum*, *P. vivax* and *P. berghei* have transmission-reducing activity. Immunisation of mice with PfHAP2 aa 195-684 induces polyclonal antibodies with comparable transmission-blocking activity to antibodies against leading transmission-blocking vaccine candidates Pfs25 and Pfs230 [15]. In addition, antibodies targeting the cd loop, a conserved putative fusion loop in PfHAP2 D2 (aa 178-195), block transmission of *P. falciparum* field isolates in a dose-dependent manner [16]. Immunisation of mice with a fusion protein comprised of PfHAP2 aa 311-609 and the cd loop also elicited transmission-reducing antibodies when formulated with various adjuvants [17]. In *P. vivax*, immunisation of PvHAP2 aa 231-459 elicited antibodies that reduce transmission [18] and in *P. berghei,* antibodies against PbHAP2 aa 355-609 [19] and the cd loop (aa 174-191) [16] reduce transmission. Previous work has also identified D3 as a region of interest, with PbHAP2 D3 (aa 477-621) capable of eliciting transmission-blocking antibodies against *P. berghei*, and showing cross-species reactivity with HAP2 of other *Plasmodium* species [8]. At present, there is no characterised monoclonal antibody against PfHAP2.

Nanobodies are the smallest naturally occurring antigen binding domain, whose small size confers advantages such as increased thermostability, high expression yields and ease of generating multivalent constructs [20]. Nanobodies are valuable tools in structural biology, acting as crystallisation chaperones that can stabilise proteins in conformational states. Transmission-reducing nanobodies have been isolated against Pfs230 [21,22] and Pfs48/45 [23], with bispecific nanobodies engineered to target both antigens showing potent transmission-reducing activity [23]. Here we aimed to isolate nanobodies targeting PfHAP2 D3 and characterise their binding and to determine the structure of PfHAP2 D3.

## Results

### Nanobodies against PfHAP2 D3 are cross-reactive with other *Plasmodium* HAP2s

To isolate nanobodies against PfHAP2, we immunised an alpaca with recombinant PfHAP2 D3 (Figure 1A and Figure S1A). We generated an immune nanobody phage display library and performed biopanning in two buffer conditions with different pH. After four rounds of biopanning, we identified 30 distinct nanobody clonal groups in which the complementarity determining region 3 (CDR3) sequence varied by three or more amino acids (WNb 321 to WNb 350) (Figure 1B, Table S1). A representative nanobody from each clonal group was selected for expression as a nanobody-Fc fusion, 22 of which successfully expressed (Figure S1B). Next-generation sequencing (NGS) of each round revealed high abundance of two closely related clones (WNb 333 and WNb 334) in rounds 3 and 4 in both buffer conditions (Figure 1C, Figure S2).

**Figure 1.**
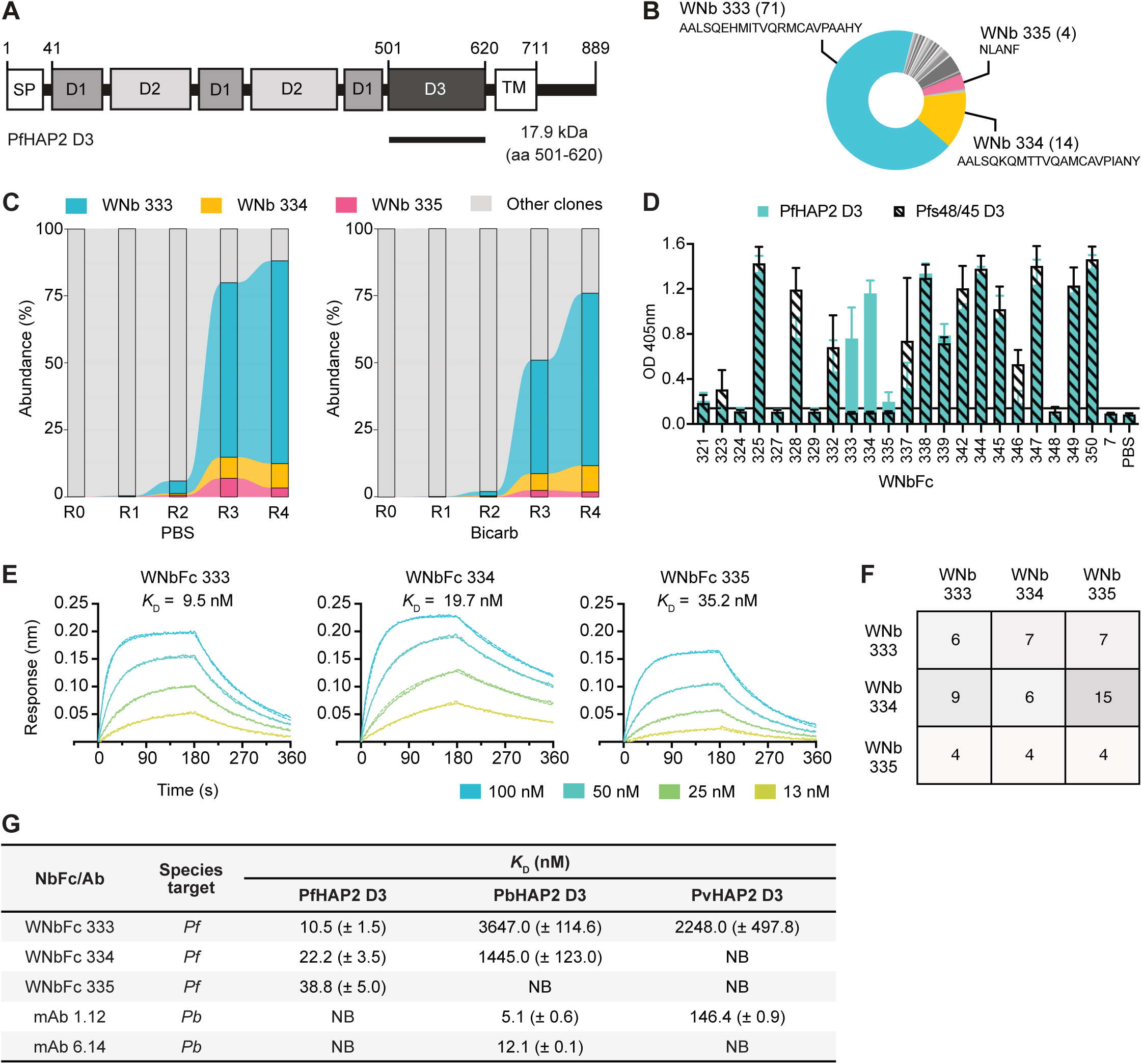
PfHAP2 D3-specific nanobodies. **A)** Schematic of full-length PfHAP2 and the recombinant construct of domain 3 (D3). Signal peptide (SP), transmembrane domain (TM) and three domains of PfHAP2 are indicated (D1, D2, D3). **B)** Overview of enrichment in the 30 anti-PfHAP2 D3 clonal groups identified by biopanning (top). PfHAP2-specific nanobodies are highlighted in colour. The number of clones per group in brackets and the CDR3 amino acid sequence are indicated. **C)** Alluvial plots showing NGS analyses of abundance of clones over four rounds of biopanning in both PBS and carbonate-bicarbonate (bicarb) buffer conditions. PfHAP2-specific clones are highlighted in colour, all other clones are coloured in grey. R0, prior to panning; R1-R4, panning rounds 1-4. **D)** ELISA of nanobody-Fcs against PfHAP2 D3 (teal) and negative control antigen Pfs48/45 D3 (hashed bars). Microtiter wells were coated with recombinant PfHAP2 D3 or Pfs48/45 D3 and probed with anti-PfHAP2 nanobody-Fcs, WNb7-Fc or PBS. Bound nanobodies were detected with HRP-conjugated anti-human IgG. Error bars represent standard deviation of the mean. **E)** Affinity curves for nanobody-Fc binding to PfHAP2 D3. Representative binding curves of two independent experiments. Different concentrations of PfHAP2 D3 to immobilised nanobody-Fcs are plotted. Binding curves were generated by bio-layer interferometry (BLI) and curves were fitted using a 1:1 Langmuir binding model. Binding affinities (*K*_D_) are indicated above binding curves. **F)** Epitope competition experiments using BLI. Nanobodies in the top row were pre-mixed with PfHAP2 D3 at a 10:1 molar ratio. Nanobodies in the left column were immobilised on the sensor and then dipped into wells with the nanobody-antigen pre-mixes. The percentage binding of the antigen-nanobody complexes was calculated relative to antigen binding alone, which was assigned to 100%. **G)** Affinities of anti-PfHAP2 nanobody-Fcs to PbHAP2 and PvHAP2 D3 by BLI. Mean affinities (*K*_D_) are given with the standard deviation if two independent experiments. *Pf*; *Plasmodium falciparum*; *Pb*; *Plasmodium berghei*; *Pv*; *Plasmodium vivax;* NB, no binding.

We tested the specificity of the nanobody-Fcs to PfHAP2 D3 using ELISA (Figure 1D). As a negative control, we used recombinant Pfs48/45 Domain 3 (Pfs48/45 D3), a *P. falciparum* 6-cysteine protein important for male fertility [24–26]. As expected, PBS and SARS-CoV-2 nanobody WNbFc 7 [27] showed no reactivity to either antigen (Figure 1D). Of the 22 nanobody-Fcs, 18 showed reactivity against PfHAP2 D3 (Figure 1D). However, the majority of these nanobody-Fcs also showed reactivity against the Pfs48/45 D3 and as such, were not specific for PfHAP2 (Figure 1D). Only three nanobody-Fcs, WNbFc 333, WNbFc 334 and WNbFc 335, showed specific reactivity for PfHAP2 D3 and not Pfs48/45 D3 (Figure 1D). We proceeded with WNbFc 333, WNbFc 334 and WNbFc 335 as they were specific to PfHAP2 D3.

Nanobody-Fc affinities for PfHAP2 D3 were determined using bio-layer interferometry (BLI) (Figure 1E, Table S2). WNbFc 333, WNbFc 334 and WNbFc 335 bound to PfHAP2 D3 with *K*_D_ values of 15.8 nM, 26.1 nM and 50.8 nM, respectively (Table S2). To determine whether these nanobodies bound to similar or distinct epitopes, we performed epitope binning by BLI. We observed that all three nanobodies competed with each other for binding to PfHAP2 D3 (Figure 1F), suggesting that the epitopes of the three nanobodies overlap.

Given anti-PbHAP2 D3 mAbs have previously shown cross-species reactivity [8], we tested our anti-PfHAP2 D3 nanobodies for reactivity against HAP2s of other *Plasmodium* species. Nanobody-Fcs and anti-PbHAP2 D3 mAbs 1.12 and 6.14 [8] were tested for binding with PfHAP2 D3, PbHAP2 D3 and PvHAP2 D3 by BLI. All nanobody-Fcs and anti-PbHAP2 D3 mAbs showed a binding response to Pf, Pb and PvHAP2 D3 above their response to the negative control Pfs48/45 D3, except WNbFc 335 and mAb 6.14, which showed no reactivity to Pv or PfHAP2 D3, respectively (Figure S3). However, we only considered cross-species reactivity where an affinity measurement could be determined. Therefore, WNbFc 333 showed cross-species reactivity against both PbHAP2 D3 and PvHAP2 D3, WNbFc 334 showed cross-species reactivity against PbHAP2 D3 only and WNbFc 335 showed no cross-species reactivity to Pb or PvHAP2 D3 (Figure 1G, Figure S3). Affinities of nanobody-Fcs for PbHAP2 D3 and PvHAP2 D3 were within the micromolar range. Anti-PbHAP2 mAb 1.12 bound to PvHAP2 D3 with a *K*_D_ value of 146.4 nM, whereas anti-PbHAP2 mAb 6.14 showed no cross-species reactivity to either Pv or PfHAP2, as previously reported [8] (Figure 1G, Figure S3).

### Crystal structure of PfHAP2 D3 in complex with nanobody WNb 334

We determined the crystal structure of PfHAP2 D3 in complex with WNb 334 to a resolution of 2.8 Å (Figure 2A, Table 1). Our refined model has a PfHAP2 D3 - WNb 334 binding interface of ∼ 640 Å, with the interaction mostly mediated by the CDR3 loop of WNb 334 (Figure 2A, right panel). Residues of the CDR3 loop form six hydrogen bonds with residues I509, I511, P512 and K616 of PfHAP2 D3 with an additional hydrogen bond formed between non-CDR residue R45 of WNb 334 and residue T507 of PfHAP2 D3 (Figure 2B, Table S3).

**Figure 2.**
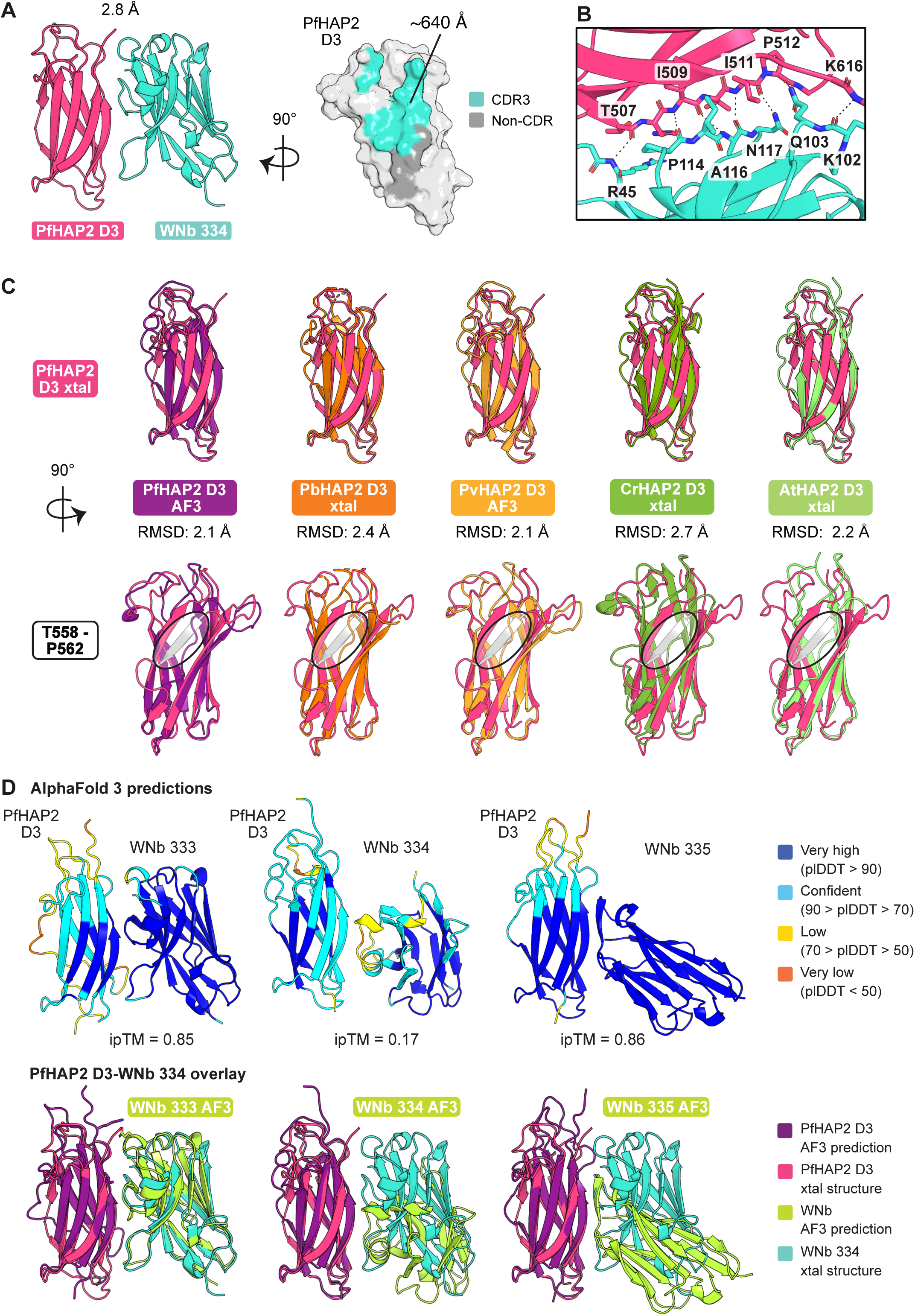
Crystal structure of PfHAP2 D3 in complex with a nanobody. **A)** Crystal structure of PfHAP2 D3 (magenta) in complex with WNb 334 (teal) at 2.8 Å (left). Surface representation of PfHAP2 D3 showing the interface with WNb 334 (right). Atoms of PfHAP2 D3 within 5 Å of CDR3 are coloured in teal, and those within 5 Å of non-CDR residues are coloured in grey. Interface surface area calculated using PISA v2.1.0. **B)** Interactions between WNb 334 (teal) and PfHAP2 D3 (magenta). Interacting residues are indicated. Hydrogen bonds are indicated by dotted line. **C)** Overlay of the PfHAP2 D3 crystal structure (magenta) with the AlphaFold 3 (AF3) PfHAP2 D3 prediction (purple), the PbHAP2 D3 crystal structure (orange, PDB 7LR4), the AF3 PvHAP2 D3 prediction (light orange), the CrHAP2 D3 crystal structure (green, PDB 6E18) and the AtHAP2 D3 crystal structure (light green, PDB 5OW3). Residues of interest (light pink) and root mean square deviation (RMSD) scores are indicated. A black circle highlights the additional β-strand (T558-P562, grey) of the PfHAP2 D3 crystal structure in the side view of the overlays. **D)** AF3 predictions of PfHAP2 D3-nanobody complexes. In the top panels, residues are coloured based on their per-residue confidence score (pLDDT). The interface predicted template modelling (ipTM) score for each prediction is indicated, representing the accuracy of the predicted relative positions of the subunits within the complex. Values higher than 0.8 represent confident high-quality predictions, while values below 0.6 suggest likely a failed prediction. Bottom panels show AF3 predictions of PfHAP2 D3 (purple) in complex with each WNb (green) overlayed with the crystal structure of PfHAP2 D3 (magenta) in complex with WNb 334 (teal).

**Table 1.**
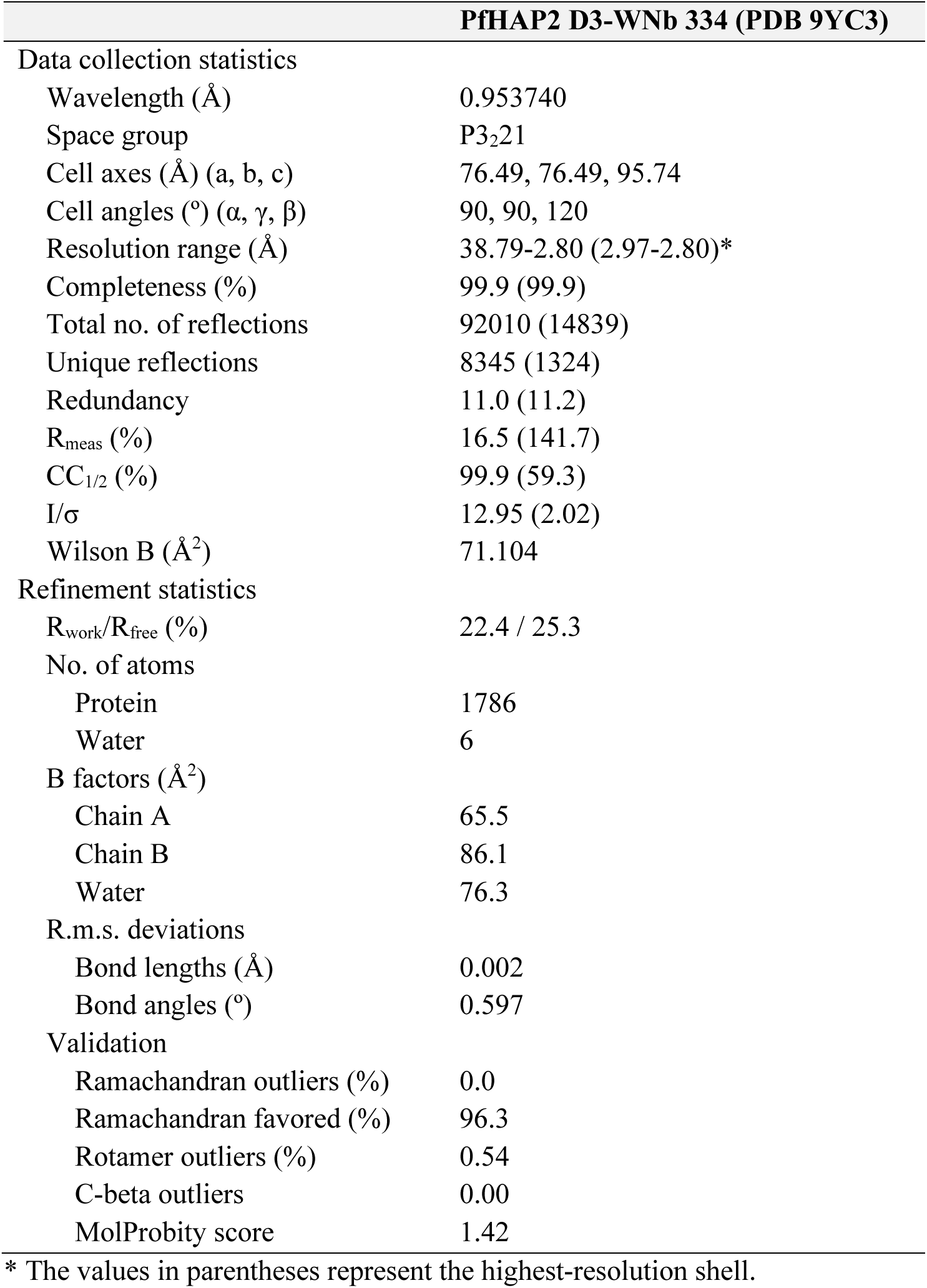
Data collection and refinement statistics for PfHAP2 D3 in complex with WNb 334.

We overlayed the PfHAP2 D3 crystal structure with its AlphaFold 3 [28] prediction with a root mean square deviation (RMSD) of 2.1 Å (Figure 2C and S4A). PfHAP2 D3 crystal structure overlays with the PbHAP2 D3 crystal structure (PDB 7LR4) [8] with an RMSD of 2.4 Å and the AlphaFold 3 prediction of PvHAP2 D3 with an RSMD of 2.1 Å. We also compared the PfHAP2 D3 crystal structure with D3 of the *C. reinhardtii* (CrHAP2) *and A. thaliana* (AtHAP2) [6,12], with RMSDs of 2.7 Å and 2.2 Å, respectively. The D3 domains of all five HAP2 proteins form a β-sandwich. Interestingly, the crystal structure of PfHAP2 D3 has eight β-strands, while the other species have seven β-strands (Figure 2C bottom panel and Figure S4B). The additional β-strand in the PfHAP2 structure (T558-P562) is not present in either the PfHAP2 D3 AlphaFold 3 prediction nor the PbHAP2 D3 crystal structure (Figure 2C, bottom panel).

With the crystal structure of the PfHAP2 D3-WNb 334 complex for comparison, we used AlphaFold 3 to predict how each of our anti-PfHAP2 D3 nanobodies would interact with PfHAP2 D3. The prediction of WNb 333 in complex with PfHAP2 D3 had high confidence, with an interface predicted template modelling (ipTM) score of 0.85 (Figure 2D, top left). Indeed, when overlayed onto the crystal structure of the PfHAP2 D3-WNb 334 complex, AlphaFold 3 predicts the WNb 333 structure and placement to be highly similar to that of WNb 334 with an RMSD of 2.9 Å (Figure 2D, bottom left). There is a high degree of sequence similarity between the two nanobodies, with only six amino acid differences among the 22 residues in their CDR3 loops (Table S1). Interestingly, of the three nanobodies, AlphaFold 3 did not predict the correct binding epitope for WNb 334 on PfHAP2 D3 (ipTM = 0.17), when compared to the experimentally validated crystal structure. We observed that the overall fold of WNb 334 of the AlphaFold prediction is highly similar with that of our crystal structure with an RMSD of 0.4 Å but the angle of approach and placement of the CDR regions towards PfHAP2 D3 differs to the PfHAP2 D3-WNb 334 crystal structure (Figure 2D, bottom middle). The prediction of WNb 335 in complex with PfHAP2 D3 was highly confident (ipTM = 0.86). The overlay suggests that WNb 335 binds a different but overlapping epitope, in agreement with our epitope binning data (Figure 2D, bottom right).

### Nanobodies against PfHAP2 D3 do not block transmission

We tested the nanobody-Fcs for transmission-blocking activity by standard membrane feeding assay (SMFA). TB31F is a potent transmission-blocking antibody against the 6-cysteine protein Pfs48/45 [29] and was used as the positive control. At 200 μg/mL, TB31F showed >99% transmission-reducing activity (TRA) across both experiments (Figure 3A). PBS and WNbFc 7, a SARS-CoV-2 specific nanobody-Fc [27] were used as negative controls, with nanobody-Fcs added to blood meals at a concentration of 100 μg/mL. The addition of WNbFc 7 did not block transmission, with only 4% TRA for both experiments (Figure 3A). None of the nanobody-Fcs showed significant transmission-blocking activity across the two experiments with averages ranging from 4 to 20% TRA across the two experiments (Figure 3A). Unfortunately, we were also unable to detect nanobody binding to activated microgametes via surface immunofluorescence assay (SIFA) (Figure 3B).

**Figure 3.**
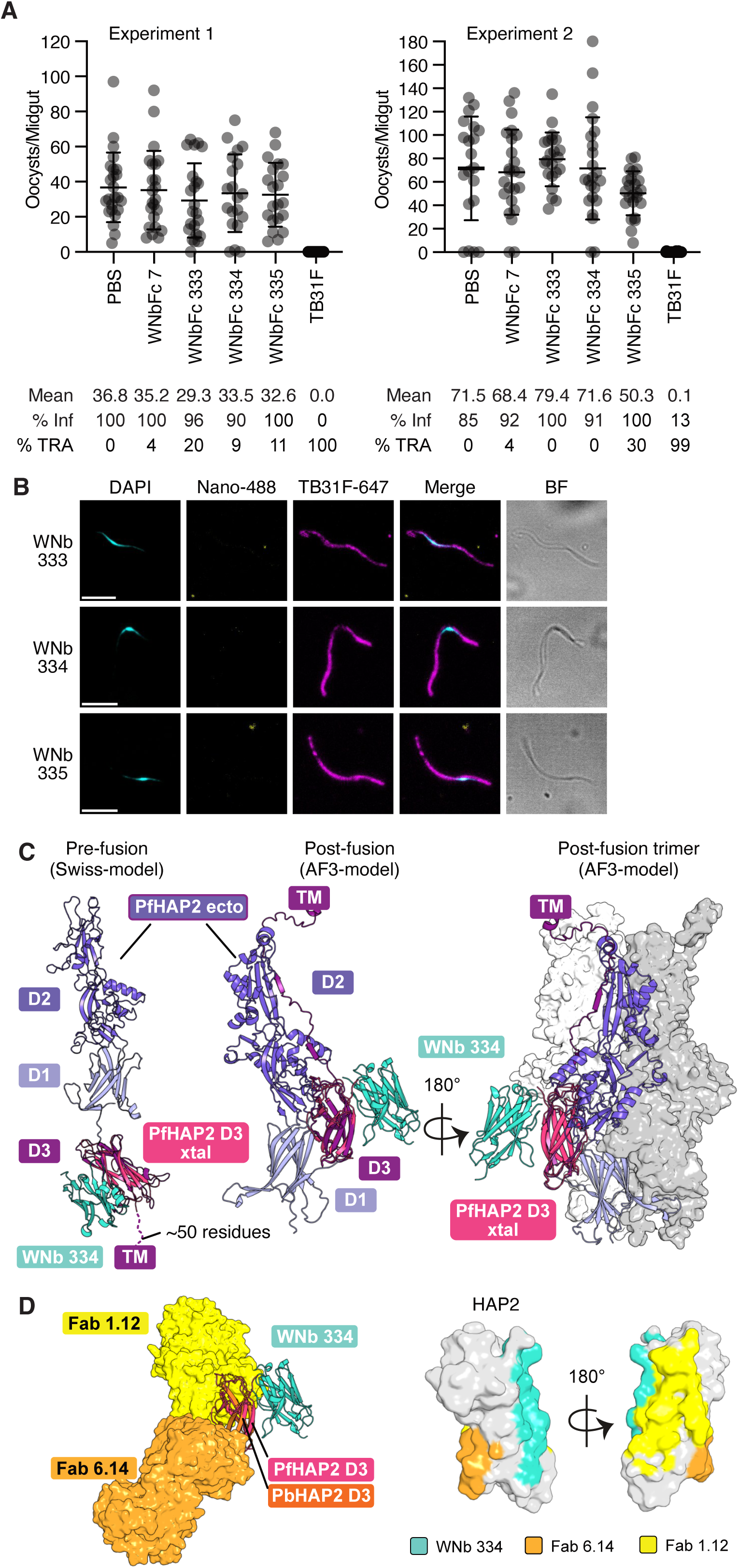
Functional characterisation of PfHAP2 D3 nanobodies. **A)** Data from two independent standard membrane feeding assay experiments testing the transmission-blocking activity of anti-PfHAP2 D3 nanobody-Fcs. *Anopheles stephensi* mosquitoes were dissected 7 - 9 days post feeding with stage V NF54 iGP2 gametocytes and oocyst numbers quantified. PBS and WNb7-Fc were included as negative controls and TB31F as a positive control. Number of oocysts per dissected midgut are plotted. Bars show means with standard deviation. % Inf = % mosquitoes infected; %TRA = transmission-reducing activity relative to the PBS control. **B)** Surface immunofluorescence assay of microgametes (activated male gametocytes) using the indicated nanobody-Fc conjugated to Alexa Fluor 488 (Nb-488, yellow) and anti-Pfs48/45 TB31F antibody conjugated to Alexa 647 (magenta). Parasite nuclei are stained with DAPI (cyan) and merged and brightfield (BF) images are shown. Scale bar = 5 μm. **C)** Crystal structure of the PfHAP2 D3-WNb 334 complex superimposed onto a pre-fusion SWISS-MODEL (based on PDB ID: 7S0K, *left*) and post-fusion AlphaFold 3 predictions (monomer: *middle*; trimer: *right*) of the PfHAP2 ectodomain (SWISS-MODEL: Y49-T614; AF3: P43-L666/R685). TM: transmembrane is indicated. **D)** Crystal structure of the PfHAP2 D3-WNb 334 complex (ribbon representation) overlayed with crystal structures of PbHAP2-Fab complexes (PDB IDs 7LR3 and 7LR4). PbHAP2 shown in orange in ribbon representation, Fab 1.12 and Fab 6.14 in surface representation, coloured yellow and light orange respectively. Middle and right panels indicate Fab and WNb 334 epitopes relative to each other on HAP2 D3 (PfHAP2 D3 used for representation).

In other organisms, HAP2 can exist as a monomer and subsequently forms a trimer during gamete membrane fusion [5,9–11]. We used AlphaFold 3 to predict the structure of the PfHAP2 ectodomain in its monomeric and trimeric forms (Figure S4A). The AlphaFold 3 prediction of the PfHAP2 monomer resembles the structures of post-fusion HAP2 ectodomains (Figure S4C) [6,12]. In addition, we generated a pre-fusion model of PfHAP2 for which we used homology modelling with SWISS-MODEL [30]. To understand how WNb 334 might bind in the context of full-length PfHAP2, we superimposed the WNb 334 binding site onto the PfHAP2 SWISS-MODEL and AlphaFold 3 predictions (Figure 3C). In both, the monomeric and trimeric structures, the WNb 334 binding site is predicted to be exposed and its binding is not predicted to block trimer formation. Using the homology with PbHAP2 D3, we also mapped the binding site of mAb 6.14 and mAb 1.12, showing that WNb 334 binds to an epitope opposite to 6.14 and adjacent to the 1.12 epitope (Figure 3D).

## Discussion

PfHAP2 is essential for malaria parasite gamete fusion during fertilisation, but its potential as a transmission-blocking candidate has not been fully explored [2]. In addition, there is no crystal structure for any domain of PfHAP2 and no monoclonal antibodies against this antigen. Here we describe a panel of nanobodies that represent the first monoclonal antibodies isolated against PfHAP2. Our nanobodies demonstrate cross-species recognition and were used to obtain the crystal structure of PfHAP2 D3, giving the first structural insights into PfHAP2.

Our PfHAP2 D3 crystal structure highlights the structural conservation of HAP2 across species, with PfHAP2 D3 showing structural similarity to HAP2 in *P. berghei*, as well as similarity across eukaryotic clades to *C. reinhardtii. and A. thaliana* HAP2 (Figure 2C). The PfHAP2 D3 crystal structure reveals an additional β-strand located from T558-P562 that is not present in the crystal structures of PbHAP2, CrHAP2, AtHAP2 or the predicted structures of PfHAP2 and PvHAP2 (Figure 2C and S4B). Therefore, the PfHAP2 D3 β-sandwich consists of two β-sheets with four β-strands each, instead of two β-sheets containing three and four β-strands as seen in other HAP2 D3 structures. This unique feature of PfHAP2 D3 will be important to consider in the design of cross-species interventions.

In many malaria-endemic areas there is concurrent transmission of both *P. falciparum* and *P. vivax* [31,32]. To achieve elimination of malaria in these regions, interventions targeting transmission of both species are required and studies of multi-species transmission-blocking vaccines are underway [33]. Recognition of HAP2 from multiple human malaria species by anti-PbHAP2 mAbs has previously been reported [8]. Here, our nanobodies against PfHAP2 show some cross-reactivity to PbHAP2 D3 and PvHAP2 D3. We observed recognition of PbHAP2 by WNb 333 and WNb 334, although there is a 294 to 418-fold reduction in affinity for PbHAP2 compared to PfHAP2 for WNb 333, and a 51 to 84-fold reduction in affinity for WNb 334 (Figure 1G, Figure S3). In the future, structure determination of our PfHAP2 nanobodies with PbHAP2 or PvHAP2 may lead to rational optimisation of cross-specific nanobodies with higher affinities, but this work is outside the scope of this paper.

We did not observe transmission-blocking activity for our nanobodies against PfHAP2 D3. There are several potential explanations for this, with various implications for the design of immunogens and prophylactics. First, all three nanobodies recognised an overlapping region of D3 that may not be essential for PfHAP2 function. A previous study by Feng *et al.* found that PbHAP2 D3 could elicit transmission-blocking antibodies against *P. berghei* [8]. Two mAbs against PbHAP2 were structurally characterised; mAb 1.12 and mAb 6.14 [8]. Both, transmission-blocking antibody mAb 6.14 and cross-reactive mAb 1.12 bind to D3 on different sites and are predicted to sterically clash with the post-fusion conformation of PbHAP2. Here, the epitope of WNb 334 is not predicted to clash with the other domains of PfHAP2 or the neighbouring chains of the trimeric post-fusion conformation (Figure 3C). It could be that mAb 6.14 and 1.12 interfere with the conformational change that HAP2 D3 undergoes during fusion, whereas WNb 334 can bind without affecting domain reorganisation and trimerisation, and therefore does not inhibit HAP2 function.

In this study, all three of our nanobodies recognised overlapping epitopes of HAP2 D3 (Figure 1F). We observed strong clonal restriction across panning rounds (Figure 1C), suggesting that nanobodies against this particular epitope are being selected for during phage display. In the future, we may be able to design nanobodies using artificial intelligence against selected epitopes such as the one bound by mAb 6.14 [34–37]. Immunisation with PfHAP2 aa 195-684 [15], the PfHAP2 cd fusion loop [16] and a fusion protein comprised of PfHAP2 aa 311-609 and the cd fusion loop [17] have all elicited antibodies with transmission-blocking activity, warranting further research into the potential of transmission-blocking nanobodies targeting these regions.

One of the major limitations of our nanobodies is their inability to recognise PfHAP2 on the surface of activated gametes in immunofluorescence-based assays. While we did observe binding of the nanobodies to recombinant PfHAP2 D3, these nanobodies may not recognize native PfHAP2 in parasites, hence the lack of detection and transmission-blocking activity. This suggests that the nanobody epitopes on recombinant PfHAP2 D3 may not be accessible or are presented differently in the pre-fusion form of PfHAP2 on the surface of malaria parasites. Structural studies on endogenous PfHAP2 using cryo-EM or crystallography will yield further insights on the folding of this critical fertilisation fusogen.

## Methods

### Recombinant protein expression and purification

The PfHAP2 D3 sequence was retrieved from PlasmoDB (PF3D7_1014200, amino acids V501-N620). DNA was codon optimised to *Spodoptera frugiperda* (*Sf*) and cloned into a modified form of baculovirus transfer vector pAcGP67-A. The PfHAP2 sequence is in frame with a GP67-signal sequence, a Tobacco Etch Virus (TEV) protease cleavage site and a C-terminal octa-histidine tag, with or without an additional AviTag. PfHAP2 D3 was produced using Sf21 cells (Life Technologies) and Insect-XPRESS Protein-free Insect Cell Medium supplemented with L-glutamine (Lonza). A cell culture of ∼1.8 × 10^6^ cells/mL was inoculated with the third passage stock of virus and incubated for three days at 28 °C. Cells were separated from the supernatant by centrifugation at 13,000 x g for 20 min. A cOmplete EDTA-free protein inhibitor tablet (Roche) and 200 mM phenylmethylsulfonyl fluoride (PMSF) was added. The supernatant was sterile filtered with a 0.45 µm filter and concentrated via tangential flow filtration using a 10 kDa molecular weight cut-off cassette (Millipore). Concentrated supernatant was sterile filtered with a 0.45 µm filter and dialysed into 30 mM Tris pH 7.5, 300 mM NaCl (buffer A). The dialysed sample was incubated with Ni-NTA resin (Qiagen) for 1 h at 4 °C on a roller shaker. A gravity flow chromatography column was washed with 10 – 20 column volumes of buffer A followed by buffer A with stepwise increases in imidazole concentration. PfHAP2 D3 eluted at imidazole concentrations of 70 mM to 200 mM. These fractions were concentrated for size exclusion chromatography (SEC) and applied to a Superdex™ 75 Increase 10/300 GL column (Cytiva) pre-equilibrated with 20 mM HEPES pH 7.5, 150 mM NaCl.

Pfs48/45 D3 (PF3D7_1346700, amino acids K293-A428) was expressed with a C-terminal octa-histidine tag and AviTag. PbHAP2 D3 (PBANKA_1212600, amino acids V499-N618) and PvHAP2 D3 (PVP01_0814300, amino acids V492-N608) were expressed with a C-terminal octa-histidine tag. Expression in Sf21 cells and purification was performed as described above for PfHAP2 D3.

Prior to alpaca immunisation and nanobody characterisation, PfHAP2 D3 HisAvi was deglycosylated with PNGase F (NEB, Cat# P0704). 1 µl of PNGase F was added per 100 µg of protein at a concentration of 2 mg/mL in 1X GlycoBuffer 2 (NEB) and incubated at 37 °C for 1-2 h. PfHAP2 D3 HisAvi was then repurified by SEC. PfHAP2 D3 HisAvi was treated with TEV protease to remove the oct-histidine and AviTag™. Protein was diluted in 20 mM HEPES pH 7.5, 150 mM NaCl and incubated with TEV overnight at 4°C. Untagged protein was separated from His-tagged TEV protease and uncleaved protein by Ni-IMAC purification using 1 mL HisTrap Excel columns (Cytiva). For crystallisation, PfHAP2 D3 His was expressed without an AviTag.

### Isolation of nanobodies against PfHAP2

One alpaca was immunised six times with ∼165 µg of recombinant PfHAP2 D3 using GERBU FAMA as adjuvant. Immunisation and handling of the alpaca for scientific purposes was approved by Agriculture Victoria, Wildlife & Small Institutions Animal Ethics Committee, project approval No. 26-17. Blood was collected three days after the last immunisation for the preparation of lymphocytes. Nanobody library construction was carried out according to established methods [21]. Briefly, alpaca lymphocyte mRNA was extracted and amplified by RT-PCR with specific primers to generate a cDNA library. The library was cloned into a pMES4 phagemid vector, amplified in TG1 *E. coli* cells and subsequently infected with M13K07 helper phage (NEB, Cat # N0315) for phage expression. Phage display was performed as previously described [21] with four rounds of biopanning on 1 µg of immobilised PfHAP2 D3. Phage display was performed with PfHAP2 D3 in either PBS or a carbonate-bicarbonate buffer (Sigma-Aldrich, Cat # C3041) at 100 mM and pH 9.5. Positive clones were identified using ELISA, sequenced using Sanger sequencing and annotated using IMGT/V-QUEST [38] and aligned in Geneious Prime 2022.1.1 (https://www.geneious.com).

### Next-generation sequencing sample preparation of nanobody phage libraries

Nanobody sequences were digested from the pMES4 phagemid vector using three different pairs of restriction enzymes to maintain sequence diversity and purified by gel extraction. Paired-end 2x300 bp sequencing libraries were prepared using the NEBNext Multiplex Oligos for Illumina (Cat # E7395) in a PCR-free manner according to the manufacturer’s instructions and sequenced on an Illumina NextSeq 2000 instrument. The NGS data was analysed using Alpseq [39]. Briefly, raw sequencing reads were trimmed to remove sequencing adapters with TrimGalore v0.6.7 (https://github.com/FelixKrueger/TrimGalore) and then merged using FLASH (v1.2.11) [40]. Annotation of nanobody sequences was performed using IgBLAST (v1.19.0) [41] with a reference database built from IMGT [42]. Nanobodies were collapsed into clones at the CDR3 level, and the counts of clones were normalised through conversion to counts per million (CPM). Alluvial plots were generated using the ggalluvial (v0.12.5) [43] package in R (v4.4.1) [44]. Cladograms were generated from generalised Levenshtein distances between CDR3s with the Neighbour-Joining method [45] from the phangorn (v2.12.1) package [46] and plotted using the ggtree (v3.14.0) R package [47]. Amino acid logos were made using the ggseqlogo R package (v0.2.0) [48].

### Nanobody and nanobody-Fc expression and purification

Nanobodies were expressed in WK6 *E. coli* cells. Bacteria were grown in Terrific Broth supplemented with 0.1% (w/v) glucose and 100 µg/mL carbenicillin at 37°C to an OD_600_ of 0.7, induced with 1 mM IPTG and grown overnight at 28°C. Cell pellets were harvested and resuspended in 20% (w/v) sucrose, 10 mM imidazole pH 7.5, 150 mM NaCl PBS and incubated on ice for 15 min. EDTA pH 8.0 was added to a final concentration of 5 mM and incubated on ice for 20 min. After this incubation, 10 mM MgCl_2_ was added, and periplasmic extracts were harvested by centrifugation at 4,000 x g for 1 h. The supernatant was sterile filtered with a 0.22 µm filter loaded onto a 1 mL HisTrap Excel column (Cytiva) in 5 mM imidazole pH 7.5, 100 mM NaCl PBS. Nanobodies were eluted with 400 mM imidazole pH 7.5, 100 mM NaCl PBS, and subsequently concentrated and buffer exchanged into 20 mM HEPES pH 7.5, 150 mM NaCl.

Nanobody sequences were subcloned into a derivative of pHLSec containing the hinge and Fc region of human IgG1 using PstI and BstEII restriction sites. Nanobody-Fc fusion proteins were expressed in Expi293 cells via transient transfection. The supernatant was harvested seven days after transfection and applied to 1 mL HiTrap PrismA affinity columns (Cytiva) equilibrated in PBS. Nanobody-Fcs were eluted in 100 mM citric acid pH 3.0 and neutralised by the addition of 1 M Tris-HCl pH 9.0. Nanobody-Fcs were subsequently buffer exchanged into PBS.

### Antibody expression and purification

The variable regions of the heavy and light chain of anti-PbHAP2 mAbs 1.12 and 6.14 [8] were synthesised as gBlocks (IDT). The heavy chain sequence was subcloned into an AbVec-hIgG1 plasmid (AddGene) using AgeI and SalI restriction enzyme sites. The light chain sequence was subcloned into an AbVec-IgKappa plasmid (AddGene), using the AgeI and HindIII restriction enzyme sites. mAbs were expressed in Expi293 cells via transient transfection using a 1:1 ratio of light and heavy chain plasmids and purified following the same protocol as described above for nanobody-Fcs. Anti-Pfs48/45 mAb TB31F [29] was expressed and purified as previously described [23].

### Purified nanobody-Fc binding ELISA

Flat-bottomed 96-well MaxiSorp plates were coated with PfHAP2 D3 or Pfs48/45 D3 diluted to a concentration of 125 nM in 50 μL PBS at room temperature (RT) for 1 h. All washes were done three times using PBS with 0.05% Tween (PBST) and all incubations were performed for 1 h at RT. Coated plates were washed and blocked by incubation with 10% skimmed milk in PBST. Plates were washed and incubated with 125 nM nanobody-Fcs in 50 μL PBS. Plates were washed and incubated with horseradish peroxidase (HRP)-conjugated goat anti-human secondary antibody (Jackson ImmunoResearch, Cat # 109-035-088) diluted 1:5000. After a final wash, 50 μL of azino-bis-3-ethylbenthiazoline-6-sulfonic acid liquid substrate (ABTS; Sigma) was added and incubated at RT for ∼20 min. 50 μL of 1% sodium dodecyl sulfate (SDS) was used to stop the reaction. Absorbance was read at 405 nm and all samples were tested in duplicate.

### BLI experiments for affinities and competition

Affinity measurements were performed on the Octet RED96e (FortéBio) using anti-human IgG Fc Capture (AHC) sensor tips (Octet®). All measurements were performed in kinetics buffer (PBS pH 7.4 supplemented with 0.1% (w/v) BSA and 0.05% (v/v) TWEEN-20) at 25°C. After a 60 s baseline step, test nanobody-Fcs or antibodies at 5 µg/mL were loaded onto sensors, followed by another 60 s baseline step. Association of nanobodies/antibodies with a two-fold dilution series of antigen was measured over 180 s, followed by dissociation in kinetics buffer for 180 s. Sensor tips were regenerated using five cycles of 5 s in 100 mM glycine pH 1.5 and 5 s in kinetics buffer. Baseline drift was corrected by subtracting the response of a nanobody-loaded sensor incubated in kinetics buffer only. Curve fitting analysis was performed with Octet Data Analysis 10.0 software using a 1:1 Langmuir binding model to determine *K*_D_ values and kinetic parameters. Curves that could not be fitted well (<0.98 R^2^ and >0.5 X^2^) were excluded from the analyses. The mean kinetic constants reported are the result of two independent experiments.

For competition BLI experiments, TEV-cleaved PfHAP2 D3 was pre-incubated with each monomeric nanobody at a 10-fold molar excess on ice for 1 h. A 30 s baseline step was established between each step of the assay. NTA sensors were first loaded with 10 µg/mL of nanobody for 5 min. The sensor surface was then quenched by dipping into 8000 nM of an irrelevant nanobody for 5 min. Sensors were then dipped into the pre-incubated solutions of PfHAP2 D3 with nanobody for five minutes. Loaded sensors were also dipped into PfHAP2 D3 alone to determine the level of PfHAP2 D3 binding to immobilised nanobody in the absence of other nanobodies. Percentage competition was calculated by dividing the premixed PfHAP2 D3 and nanobody solution binding at 100 seconds by the PfHAP2 D3 binding at 100 seconds alone, multiplied by 100.

### Crystallisation and structure determination

PfHAP2 D3 and nanobody WNb 334 were complexed at a molar ratio of 1:1.4 for 1 h on ice. Complexes were applied to a Superdex™ 75 Increase 10/300 GL column (Cytiva) pre-equilibrated in 20 mM HEPES pH 7.5, 150 mM NaCl and purified by SEC. Sitting drop initial and optimisation screens of PfHAP2 D3-WNb 334 were set up at the Monash Macromolecular Crystallization Platform (MMCP, Clayton, VIC, Australia). Initial crystals of complex at 11.2 mg/mL grew in 0.1 M HEPES pH 7.5, 25% w/v PEG 3350 at 20 °C and were optimised by increasing the concentration of protein to 15 mg/mL, the pH of HEPES to 7.7, and concentration of PEG 3350 to 27%. Crystals were harvested with 30% MPD in mother liquor before flash freezing in liquid nitrogen.

X-ray diffraction data was collected at the MX2 beamline of the Australian Synchrotron [49]. The XDS package [50] was used for data processing. Molecular replacement was used to solve the phase problem using the AlphaFold 2 (v.2.3.2) [51] predictions of PfHAP2 D3 and nanobody WNb 334 structures. Iterative cycles of structure building and refinement were carried out using Coot v 0.9.8.92 [52] and Phenix v 1.21-5184 [53,54]. Interface surface area and bonds were determined by PISA v 2.1.0 [55]. Figures of the structure were prepared with PyMOL v 2.5.0 [56]. Structural alignments were performed using *super* in PyMOL and all-atom RMSD values are reported. The atomic coordinates and structure factor files have been deposited in the Protein Data Bank under PDB ID 9YC3.

### *P. falciparum* maintenance and gametocyte culture

*P. falciparum* NF54 iGP2 [57] asexual parasites were cultured in human type O+ erythrocytes (Melbourne Red Cross LifeBlood) at 4% haematocrit in RPMI HEPES supplemented with 50 µg/mL hypoxanthine, 0.2% (w/v) NaHCO_3_, and 10 mM D-Glucose containing 5% heat-inactivated human serum and 5% Albumax (ThermoFisher Scientific). iGP2 gametocytes for transmission to mosquitoes were generated as previously described [57]. Gametocyte cultures were maintained in RPMI 1640 medium supplemented with 26 mM HEPES, 50 µg/mL hypoxanthine, 0.2% (w/v) NaHCO_3_, and 10 mM D-Glucose, containing 10% heat-inactivated human serum and with daily media changes. Use of human blood and serum for parasite culturing was approved by the WEHI Human Research Ethics Committee, project approval No. 86-17.

### Surface immunofluorescence assay of *P. falciparum* activated gametes

Stage V gametocytes were harvested by spinning at 13,000 rpm for 1 min. Gamete activation was induced by resuspending in 400 µl ookinete medium (RPMI HEPES with 100 uM xanthurenic acid) containing 10 µg/mL anti-Pfs48/45 TB31F directly conjugated to Alexa Fluor 647 and 10 µg/mL anti-PfHAP2 nanobody-Fc directly conjugated to Alexa Fluor 488. Antibody/nanobody conjugation was carried out by the WEHI Antibody Facility. The suspensions were incubated on 0.1 mg/mL PHA-E treated (Sigma) glass coverslips in humidified chamber at room temperature for 30 min. Slides were washed three times with 1X PBS and then activated gametes were fixed with 2% formaldehyde in PBS for 20 min in a humidified chamber. The slides washed again and mounted with a coverslip using VECTASHIELD PLUS antifade mounting medium with DAPI (Vector Laboratories). Coverslips were sealed with nail polish. Images were acquired on a Zeiss LSM 980 using confocal mode with a 63x (1.4NA) objective lens with oil immersion using the 405, 488 and 561 lasers. Images were processed using FIJI ImageJ software (v2.14.0/1.54) [58] and the Z-projection max intensity images were used for figures.

### Standard membrane feeding assays

Three- to five-day old female *Anopheles stephensi* mosquitoes were fed via a water-jacketed glass membrane feeder. For nanobody-Fc experiments, blood meals were comprised of 250 µL O+ heat-inactivated human serum and 250 µL O+ erythrocytes with a stage V gametocytemia of ∼0.4%. To blood meals, 10 µL nanobody-Fcs at 5 mg/mL in PBS, 10 µL TB31F at 10 mg/mL in PBS or 10 µL PBS was added. Blood-feeding was allowed to proceed for 1-2 h and engorged mosquitoes were housed in cups and provided with sugar water. After 7-8 days, midguts were dissected from cold-anaesthetised and ethanol sacrificed mosquitos. Midguts were stained with 0.1% mercurochrome and oocyst numbers were counted. Data were plotted and statistical analyses were performed using GraphPad Prism 9 version 9.5.1 for Mac, GraphPad Software, Boston, Massachusetts USA, www.graphpad.com. Statistical significance was determined using one-way ANOVA and Tukey’s multiple comparison test. **** P ≤ 0.0001. *** P ≤ 0.001. ns P > 0.05.

## Data availability

Coordinates and structure factor files of the PfHAP2 D3-WNb 334 crystal structure have been deposited in the Protein Data Bank under PDB ID 9YC3 [59]. The NGS data has been deposited to the European Nucleotide Archive (ENA) under accession number PRJEB101643 [60].

## Acknowledgments

We thank Geoffrey Kong from the Monash Macromolecular Crystallisation Platform (MMCP, Clayton, VIC, Australia) for assistance with setting up the crystallisation screens. This research was undertaken using the MX2 beamline at the Australian Synchrotron, part of ANSTO, and made use of the Australian Cancer Research Foundation (ACRF) detector. We thank the MX2 beamline staff at the Australian Synchrotron for their assistance during data collection. NF54/iGP2 was kindly provided by Till Voss (Swiss Tropical and Public Health Institute). W-H.T. is supported by National Health and Medical Research Council of Australia (NHMRC) GNT2016908 and APP2001385. Q.G. is supported by NHMRC GNT2007996. We thank the Australian Red Cross Lifeblood for the supply of human erythrocytes and serum. The authors acknowledge the Victorian State Government Operational Infrastructure Support and Australian Government NHMRC IRIISS.

## Supplementary material

**Figure S1.**
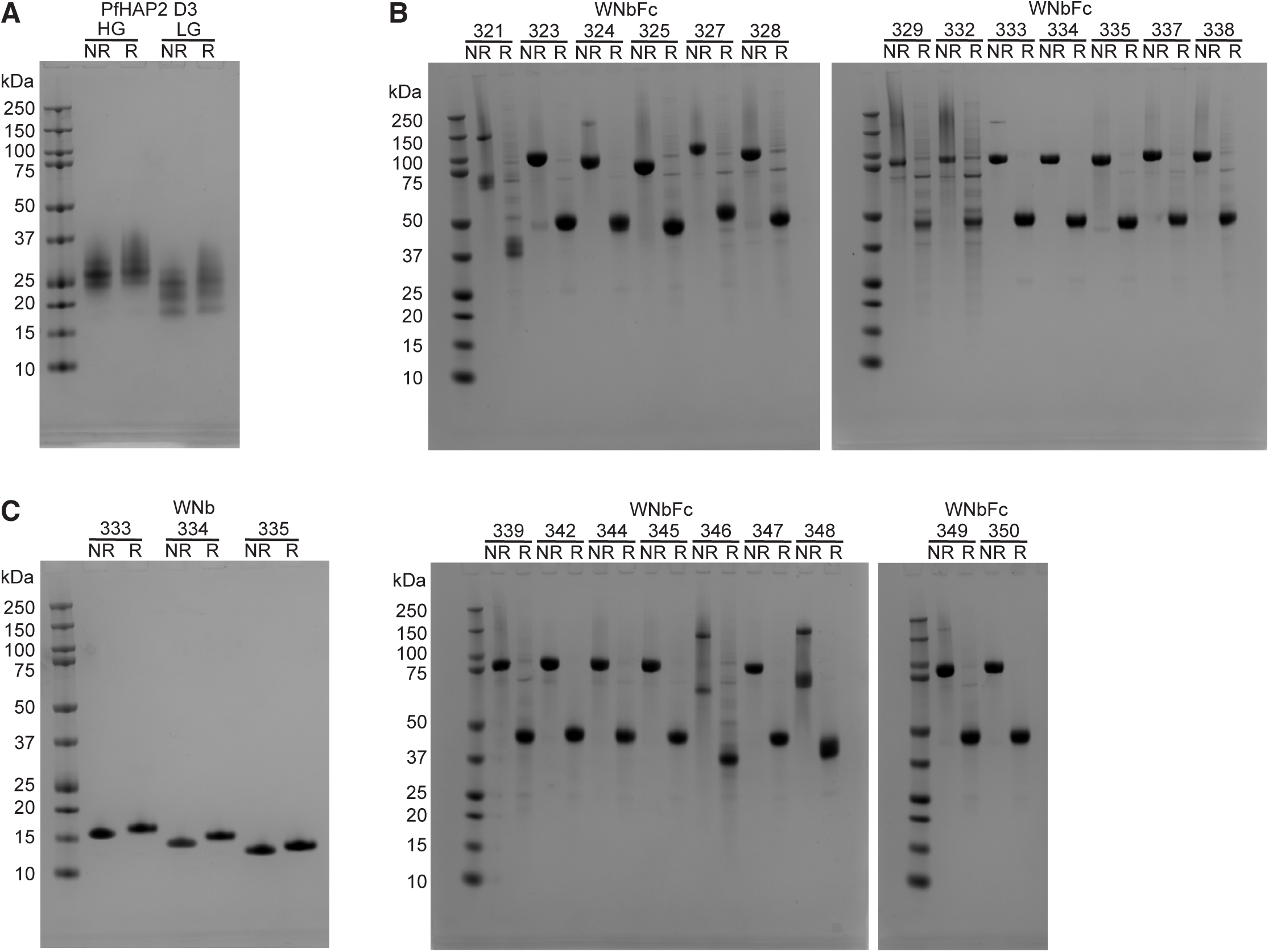
Recombinant protein expression. **A)** SDS-PAGE of purified PfHAP2 D3 antigen under non-reducing (NR) and reducing (R) conditions. Highly glycosylated (HG) and less glycosylated (LG) fractions are indicated. **B)** Purified PfHAP2 nanobody-Fcs under non-reducing and reducing conditions. **C)** Purified PfHAP2 nanobodies under non-reducing and reducing conditions.

**Figure S2.**
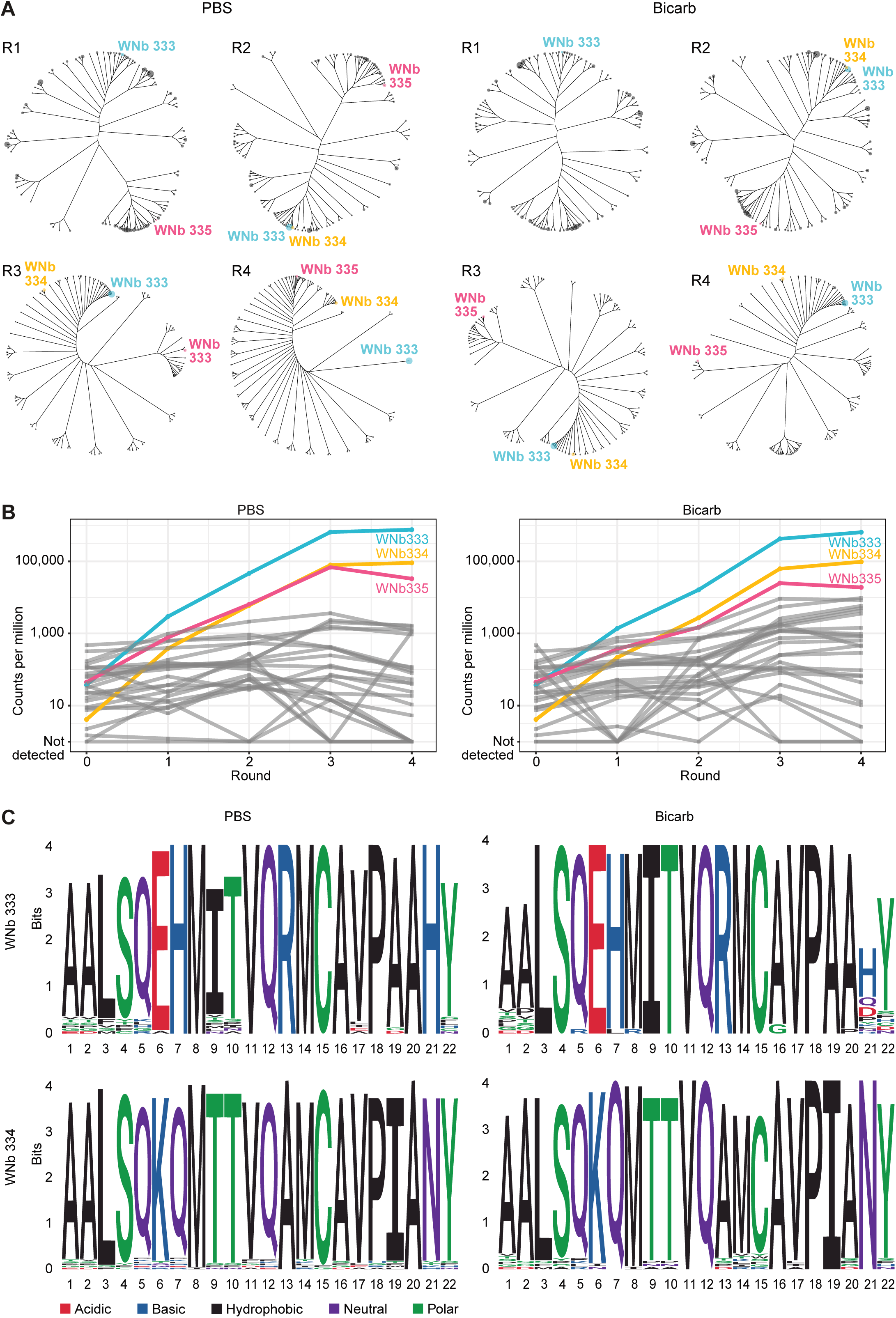
Next-generation sequencing (NGS) analysis of panning libraries. **A)** Cladogram of the 100 most abundant nanobodies from NGS of rounds 1 - 4 phage display selection libraries. PfHAP2-specific clones are labelled. Tree tips are scaled relative to abundance (counts per million). **B)** Line graph of the normalised counts per million (CPM) from next-generation sequencing analysis of the clones identified by Sanger sequencing before panning (round 0) and across rounds 1-4. PfHAP2-specific clones are labelled. **C)** Sequence logo plot showing the sequence conservation of amino acids in the complementarity-determining region 3 (CDR3) loop of clones within the WNb 333 and WNb 334 clusters. Letter height is proportional to residue conservation within the cluster.

**Figure S3.**
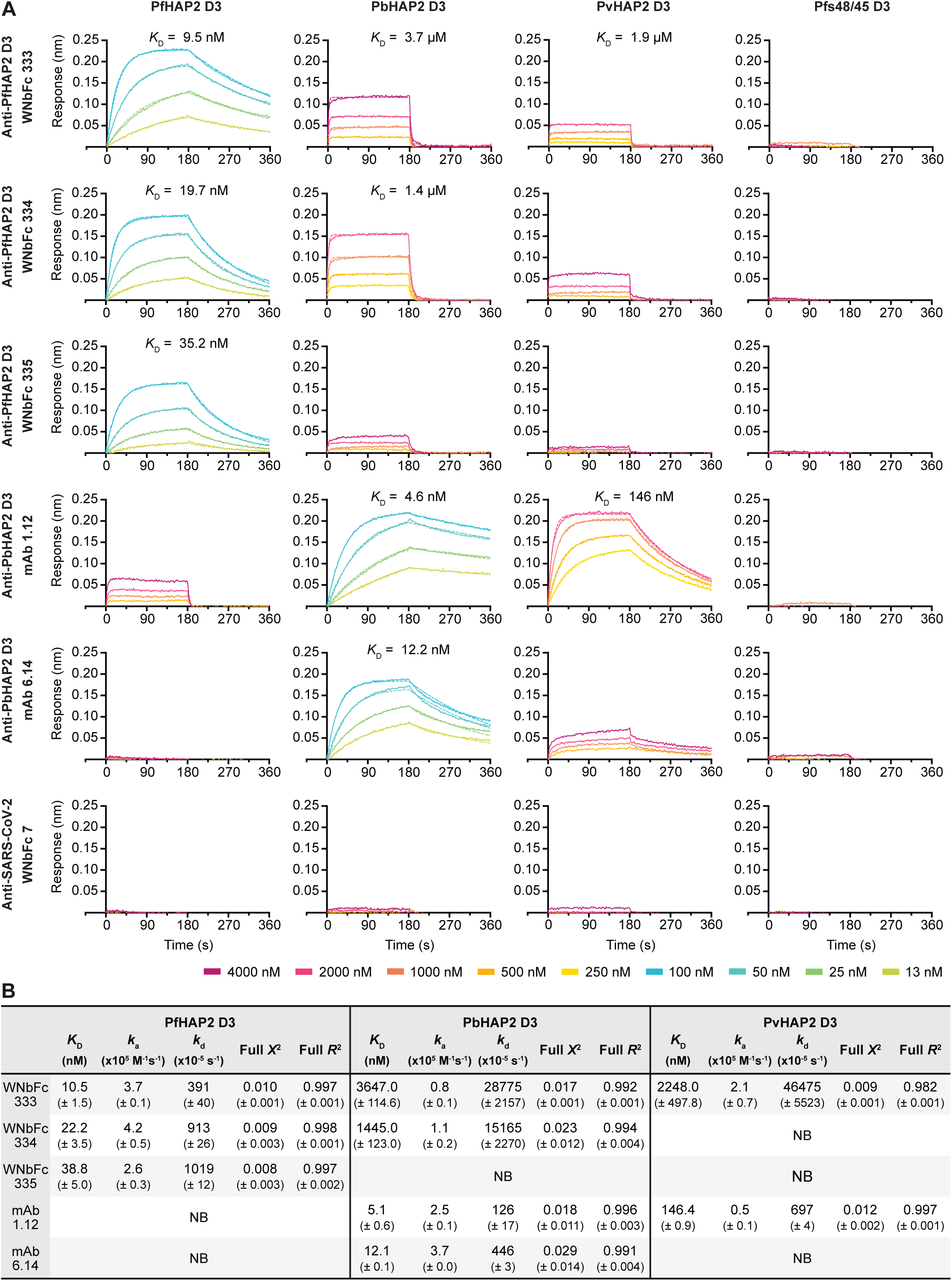
Cross-reactivity of anti-PfHAP2 nanobody-Fcs with HAP2 of different *Plasmodium* species. **A)** Representative binding curves of different concentrations of antigen to immobilised nanobody-Fcs or mAbs. Binding curves were generated by bio-layer interferometry (BLI) and curves were fitted using a 1:1 Langmuir binding model. Binding affinities (*K*_D_) are indicated above binding curves. **B)** Affinities of anti-PfHAP2 nanobody-Fcs to PbHAP2 and PvHAP2 D3 by BLI. Mean affinities (*K*_D_), association rates (*k*_a_), dissociation rates (*k*_d_) are given with the standard deviation of two independent experiments. *Pf*; *Plasmodium falciparum*; *Pb*; *Plasmodium berghei*; *Pv*; *Plasmodium vivax;* NB, no binding.

**Figure S4.**
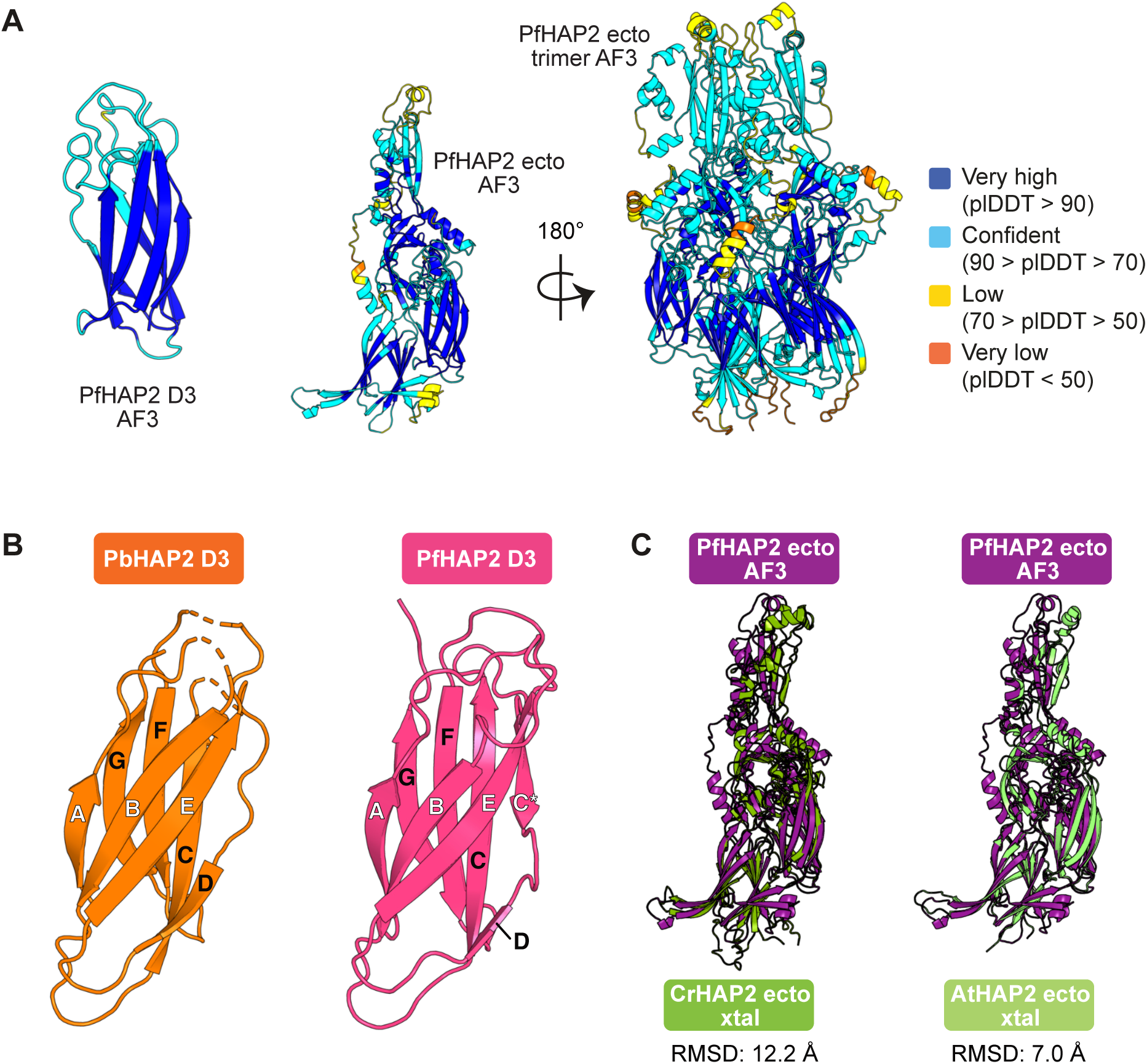
AlphaFold 3 (AF3) prediction of PfHAP2 structures. **A)** AF3 predictions of PfHAP2 D3, full-length and trimer structures. The full-length and trimer structures were truncated to display the ectodomain only (P43-V621). Residues are coloured based on their per-residue confidence score (pLDDT). **B)** Left: PbHAP2 D3 (PDB ID: 7LR4) contains seven β-strands (labelled A to E) that form two β-sheets, of which one contains β-strands ABE and the other DCFG. Right: PfHAP2 D3 contains eight β-strands. In comparison to PbHAP2 D3 it has an additional β-strand, labelled C*, which is located between β-strands C and D. Its β-sandwich is formed by two β-sheets containing the β-strands ABEC* and DCFG. **C)** Overlay of the AF3 prediction of the PfHAP2 ectodomain structure (purple) with the determined crystal structures for the CrHAP2 ectodomain (green, PDB 6E18) and AtHAP2 ectodomain (light green, PDB 5OW3). Root mean square deviation (RMSD) scores are indicated.

**Table S1.**
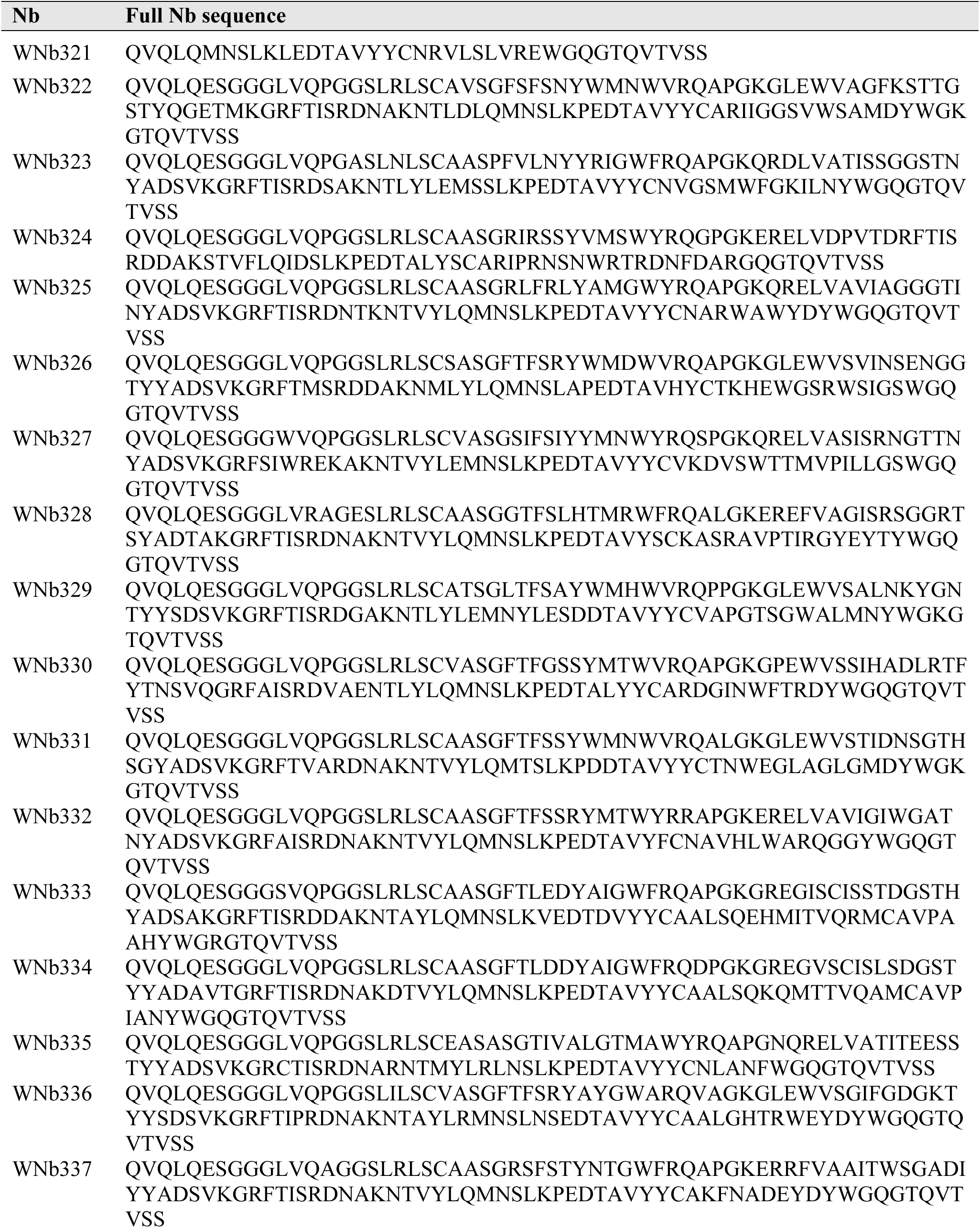
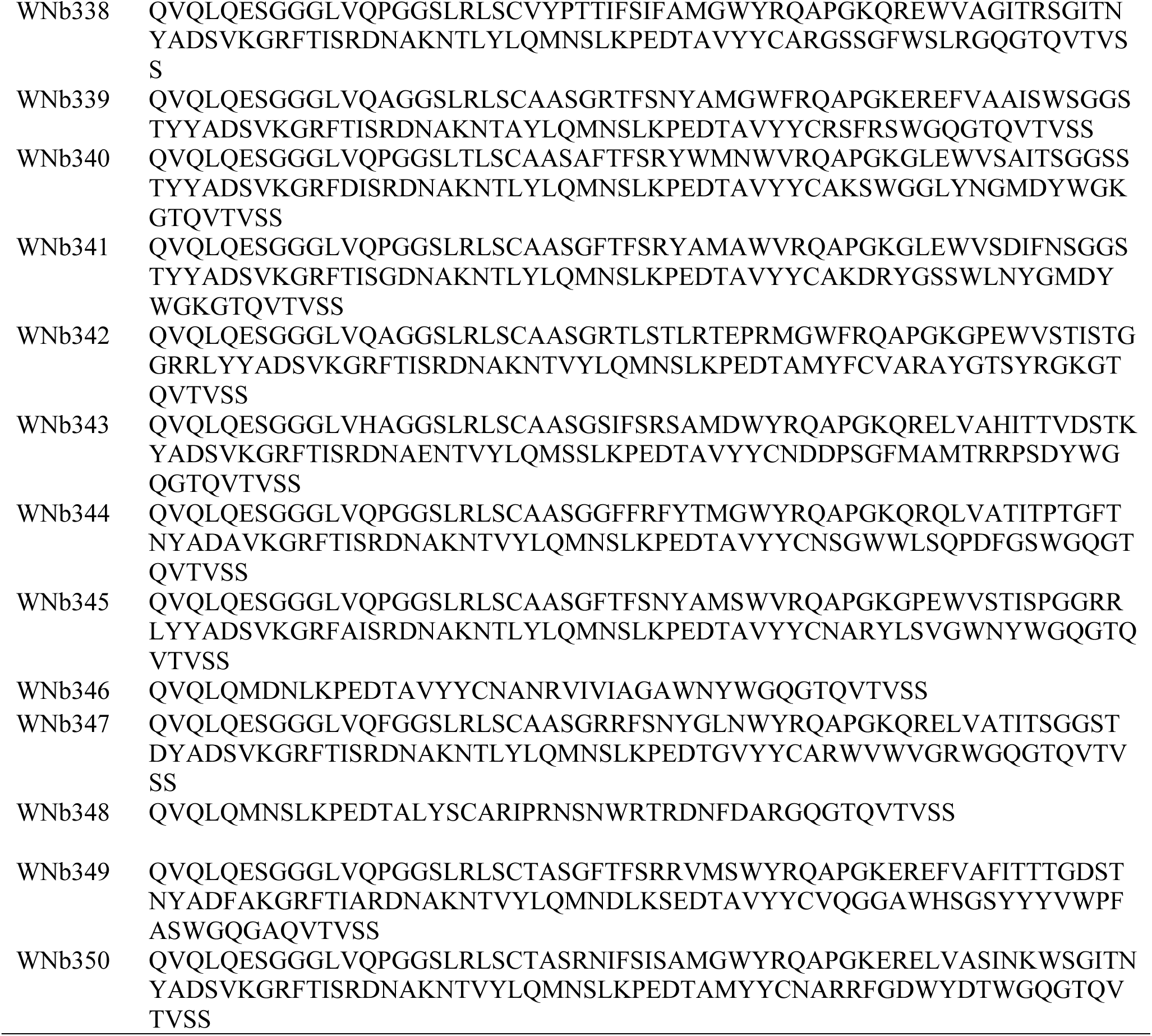
Amino acid sequences of nanobodies (Nb) isolated against PfHAP2 D3.

**Table S2.** Affinities of nanobody-Fcs for PfHAP2 D3 by bio-layer interferometry (BLI).

| | $K_D$ (nM) | $k_a$ ($\times 10^5$ M<sup>-1</sup>s<sup>-1</sup>) | $k_d$ ($\times 10^{-5}$ s<sup>-1</sup>) | Full $X^2$ | Full $R^2$ |
| --- | --- | --- | --- | --- | --- |
| WNb333 | 15.8 ± 1.3 | 3.6 ± 0.5 | 567.8 ± 35.7 | 0.008 ± 0.003 | 0.996 ± 0.001 |
| WNb334 | 26.1 ± 0.1 | 4.4 ± 0.4 | 1144.5 ± 105.5 | 0.013 ± 0.001 | 0.995 ± 0.001 |
| WNb335 | 50.8 ± 0.9 | 1.9 ± 0.2 | 990.6 ± 121.4 | 0.006 ± 0.001 | 0.996 ± 0.001 |

Mean affinities ( $K_D$ ), association rates ( $k_a$ ), dissociation rates ( $k_d$ ), $X^2$ and $R^2$ values of two independent experiments are given with standard deviation.

**Table S3.** Summary of interactions between PfHAP2 D3 and WNb 334 using PISA.

| PfHAP2 D3 | Group | WNb 334 | Group | Distance (Å) |
| --- | --- | --- | --- | --- |
| *Hydrogen bonds* |  |  |  |  |
| Thr 507 | O | Arg 45 | N | 3.1 |
| Ile 509 | N | Pro 114 | O | 3.1 |
| Ile 509 | O | Ala 116 | N | 3.4 |
| Ile 511 | N | Ala 116 | O | 3.0 |
| Ile 511 | O | Asn 117 | ND2 | 3.0 |
| Pro 512 | O | Gln 103 | NE2 | 3.0 |
| Pro 512 | O | Asn 117 | ND2 | 3.4 |
| Asn 534 | ND2 | Gly 42 | O | 3.7 |
| Lys 616 | N | Lys 102 | O | 3.5 |
| Other WNb 334 interfacing residues (PfHAP2 D3) |  |  |  |  |
| Ser 502 | Tyr 503 | Thr 505 | Ile 506 | His 508 |
| Thr 510 | Lys 513 | Cys 515 | His 530 | Trp 532 |
| Leu 609 | Ser 610 | Phe 611 | Asn 612 | Leu 613 |
| Thr 614 | Ser 615 |  |  |  |
| Other PfHAP2 D3 interfacing residues (WNb 334) |  |  |  |  |
| Tyr 32 | Gln 39 | Pro 41 | Lys 43 | Gly 44 |
| Leu 99 | Ser 100 | Thr 105 | Val 113 | Ile 115 |
| Tyr 118 | Trp 119 |  |  |  |

